# CLASPP: A unified model for predicting post-translational modifications

**DOI:** 10.64898/2026.06.04.729962

**Authors:** Nathan Gravel, Zhongliang Zhou, Ruili Fang, Austin Downes, Saber Soleymani, Natarajan Kannan

**Affiliations:** Institute of Bioinformatics, University of Georgia, GA 30602, USA; School of Computing, University of Georgia, GA 30602, USA; Institute for Insight, Georgia State University, Atlanta, GA 30303, USA; Department of Biochemistry and Molecular Biology, University of Georgia, GA 30602, USA

## Abstract

Post-Translational Modifications (PTMs) are a fundamental mechanism for regulating cellular pathways and increasing the functional diversity of the proteome. Accurately predicting the PTM types that are likely to occur at a given site in the primary sequence is a key challenge in functional proteomics. Existing PTM prediction models predominantly focus on either single PTM types or employ ensemble methods that combine multiple models to predict different PTM types. This fragmentation is largely driven by the vast imbalance in data availability across PTM types, making it difficult to predict multiple PTM types with a single model. To address this limitation, we present the **C**ontrastively **L**earned **A**ttention-based **S**tratified **P**TM **P**redictor (CLASPP), a unified PTM prediction model. CLASPP addresses imbalance challenges by leveraging unsupervised clustering-based undersampling and a novel contrastive learning framework tailored to PTM data. Additionally, our hierarchical data organization and curation are shown to improve CLASPP’s performance by balancing the representation of individual PTM types and provides a standardized dataset to train and validate future model designs. Drawing inspiration from advancements in image and natural language processing, the CLASPP model employs a multi-stage training strategy and a high-quality, curated training dataset to improve PTM prediction performance. To uncover what is learned during the contrastive learning stage, the CLASPP model is shown to distinguish known protein kinase substrate specificity profiles as a form of explainability. Finally, we evaluate the application of CLASPP in predicting PTMs in different model organisms and experimentally validated ubiquitination sites in the understudied DCLK3 kinase. Overall, CLASPP represents a unified model for PTM prediction that addresses key bottlenecks in data imbalance and offers new strategies for biological data curation, thereby improving PTM-type prediction performance across diverse organisms.

**Author summary:** Post-translational modifications (PTMs) are essential changes that proteins undergo, influencing nearly every aspect of cell function, communication, and disease. Accurately predicting where and how these modifications occur is challenging due to the diversity of PTM types and the limitations of existing annotation pipelines. This study introduces a unified deep learning approach, termed CLASPP, leveraging contrastive learning to predict multiple PTM types simultaneously from primary protein sequence alone. By employing advanced data balancing and sampling methods, CLASPP ensures reliable predictions for rare and common modifications. The model utilizes a pre-trained protein language model to capture sequence and structural features encoded in the primary protein sequence. Test results demonstrate that CLASPP consistently surpasses existing tools in predicting 12 major PTM types, and its versatility enables robust predictions across species, not just in human proteins. The final model, data curation, and training datasets are freely accessible for broader use and reproducibility (https://github.com/gravelCompBio/Claspp_forward).

## Introduction

Post-Translational Modifications (PTMs) play a pivotal role in regulating cellular functions and enhancing the functional diversity of proteins across all domains of life [1, 2]. Current data indicates that approximately 78% of the human proteome is associated with at least one PTM, as documented in the neXtProt database [3, 4]. While well-known modifications like Serine/Threonine/Tyrosine phosphorylation are extensively studied due to their crucial roles in cellular function [5–7], many other PTM types remain largely unexplored [1, 8]. Despite advances in Mass-Spectrometry (MS) based detection methods and the cataloguing of multiple PTM types in databases such as dbPTM [9], PhosphoSite+ [10], UniProt [11], and neXtProt [3], challenges persist in detecting and annotating multi-PTM events [12]. Specifically, when multiple amino acids within a peptide are modified, the resulting combinations dramatically increase the complexity of the search space, posing a significant computational challenge for annotating multiple PTM events [13]. Furthermore, the dynamic nature of PTMs and the resulting fragmentation patterns during tandem mass spectrometry makes identification and annotation of many PTM types difficult using experimental approaches alone [14, 15].

The uneven distribution of PTM data further complicates the prediction process, as current datasets are skewed towards a subset of well-studied PTMs types [8, 16]. PTM types such as phosphorylation, ubiquitination, acetylation, glycosylation, and methylation are generally overrepresented in existing databases, creating an imbalance in training datasets for downstream prediction algorithms [16, 17]. Traditionally, the data imbalance bottleneck has been addressed by applying a sequence identity filter [18–21] which, while helpful, fails to directly address the long-tail distribution in which a few PTM types are overrepresented relative to others. To overcome this long-tail distribution problem, Zuo et al. (2024) demonstrated that k-means clustering based undersampling of sequence features increases performance towards predicting multiple lysine modifications using a convolutional neural network [22]. Likewise, data stratification strategies have been shown to improve model performance in other bioinformatics applications such as cancer subtype classification [23] and protein engineering [24]. Although, strategies to overcome data imbalance issues in a multi-PTM prediction task have been proposed [25], they have not been systematically evaluated.

Recent advances in deep learning, particularly in natural language processing, have introduced the use of masked language modeling on protein sequences, a self-supervised training method that enables unbiased extraction of evolutionary [26, 27], structural [28–30], and functional characteristics [31, 32] from a vast amount of protein sequence data [33]. More recently, protein language models [34, 35] have achieved notable success by incorporating contrastive learning with traditional mask language modeling [36, 37]. Originally developed for image classification, contrastive learning has been successfully employed for a broad range of biological applications [38]. Contrastive learning encourages models to more effectively distinguish between similar and dissimilar representations by bringing similar objects/data points closer to each other in the latent variable space while pushing dissimilar data points farther apart [39]. Indeed, contrastive loss functions have been shown to improve model performance in biological function prediction tasks [40, 41], sequence classification [42], and annotation of rare PTMs [43]. Nonetheless, the impact of contrastive learning in enhancing multi-PTM prediction performance has yet to be comprehensively assessed.

Existing PTM prediction models generally fall into two main categories: specific and multi-PTM prediction approaches. Specific PTM models focus exclusively on predicting a single PTM type such as Emeber [44], UbiSite [45], DEEPPRMS [46], TransPTM [47], and Stack-OglyPred-PLM [48]. On the other hand, multi-PTM models are designed to predict multiple PTM types simultaneously, often adopting an ensemble modeling approach. Examples of multi-PTM prediction models include Musitedeep [18], MIND-S [19], PTMGPT2 [20], AstraPTM [49] and Sitetack [50]. Additionally, other PTM-centric deep learning models of note includes, DeepMVP [21], MTPrompt-PTM [25], Phosformer-ST [32, 51], COMPASS-PTM [52] and PTM-mamba [53]. DeepMVP was trained on datasets that are validated with multiple sources to ensure a high-quality training set [21]. MTPrompt-PTM is a fine-tuned version of S-PLM which was originally trained on protein sequence and 3D structural data [54]. Phosformer-ST leverages the Serine/Threonine Substrate Specificity Atlas dataset [55] to train the model with an added input of PTM transferase (protein kinase) sequences, unlike most other models that rely solely on the modified substrate sequence as input. Additionally, COMPASS-PTM builds on this framework of enzyme specific PTM prediction and predicts the enzyme involvement for multiple PTM types. PTM-mamba [53] introduces a generative model that was trained with PTM data along with traditional protein sequences to improve generative and protein engineering tasks. Thus, although multiple PTM prediction models have been proposed, they have not been systematically benchmarked to inform future development efforts.

In this work, we systematically benchmark existing multi-PTM prediction models and introduce **C**ontrastively **L**earned **A**ttention-based **S**tratified **P**TM **P**redictor (CLASPP), a unified model designed to address data imbalances and improve the performance of multi-PTM predictions. CLASPP employs a three-stage training regimen integrating masked language modeling [56], contrastive learning [39], and finetuning [30, 56, 57], alongside an optimized unsupervised clustering [58, 59] based data curation. CLASPP is tested with multiple encoders of different protein language model architectures [30, 60, 61] and demonstrates superior global performance over other multi-PTM prediction models. It offers explainability by separating phosphorylated peptide sequences in the contrastive learning step, which resembles known kinase substrate specificity profiles [55]. Finally, CLASPP’s practical utility is demonstrated through out-of-distribution performance on PTM types from different model organisms and the prediction of experimentally validated ubiquitination sites on the understudied protein kinase DCLK3 (UniProt ID : Q9C098) [62, 63].

## Results

### Contrastively Learned Attention-based Stratified PTM Predictor (CLASPP) model design and training

The CLASPP model was built upon an ESM-2-150m encoder, integrated with a multi-binary classification head to enable simultaneous prediction across multiple PTM types (Fig 1a). This unified approach begins with the encoder undergoing pre-training using a masked language modeling task, leveraging protein sequences from the UniRef database to establish residue-specific relationships among millions of proteins [33] (Fig 1b). Following this, the encoder was refined through a supervised contrastive learning stage [64] utilizing a stratified PTM dataset, designed to better separate representation among the 12 PTM types (Fig 1b). The final stage involved finetuning the classification head on the PTM prediction tasks, during which the encoder remained frozen, preserving the learned features from the previous contrastive learning stage (Fig 1b).

**Fig 1.**
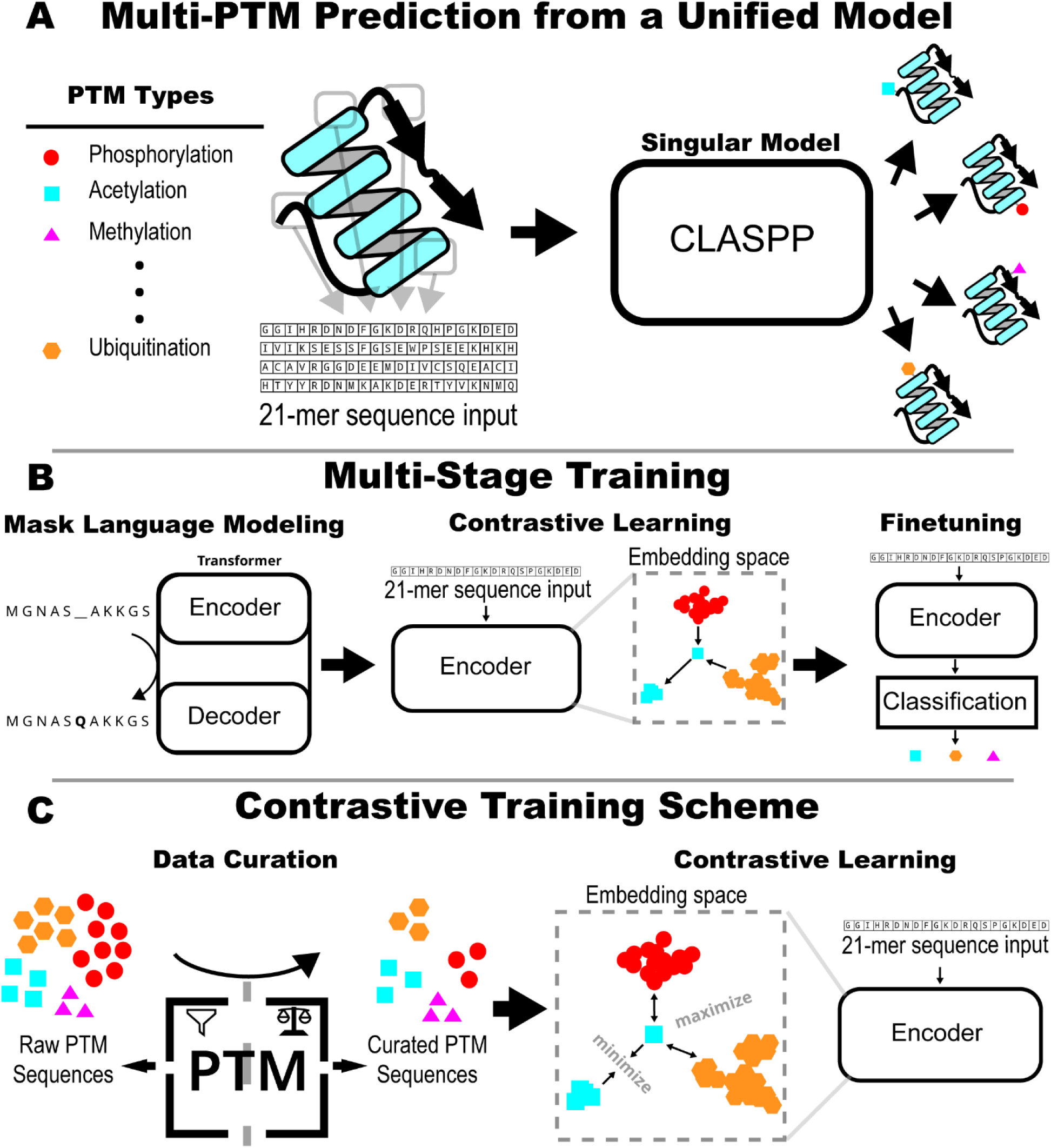
Overview of the CLASPP model. (A) Demonstrates the multi-PTM prediction capability of the unified CLASPP model. Each inference of the model takes a 21-mer peptide with center residue being the modified residue. (B) Shows the 3-stage training of the CLASPP model. (C) Highlights the major innovation. Incorporating improved data curation and stratification into a contrastive learning framework.

The innovation of CLASPP lies in its multi-stage training strategy, particularly the integration of custom hierarchical data into the supervised contrastive learning phase (Fig 1c). Comparative evaluations include alternative architectures, such as ESM-C-300m, ESM-C-600m, ESM-3-open [60, 61], and a multi-headed attention classification head (S1 Fig), highlighting the improvements in CLASPP’s unified framework [49]. Notably, unlike ensemble methods that stitch together multiple models [18–21, 25], CLASPP offers a streamlined, single-model solution for multi-PTM prediction [43, 49]. The final model comprises 1.4 billion parameters.

### Addressing data imbalance and optimizing data clustering enhances prediction performance

Uneven representation among PTM types poses a significant challenge for multi-PTM prediction performance. To tackle this issue, we employed a systematic PTM data curation process (Fig 2a, b, and c; see Methods; see S2 Fig), which includes unsupervised clustering combined with uniform sampling. By deriving sub-clusters for each PTM type and applying uniform sampling to each PTM sub-cluster, we observed a notable improvement in downstream PTM prediction performance (Fig 2d).

**Fig 2.**
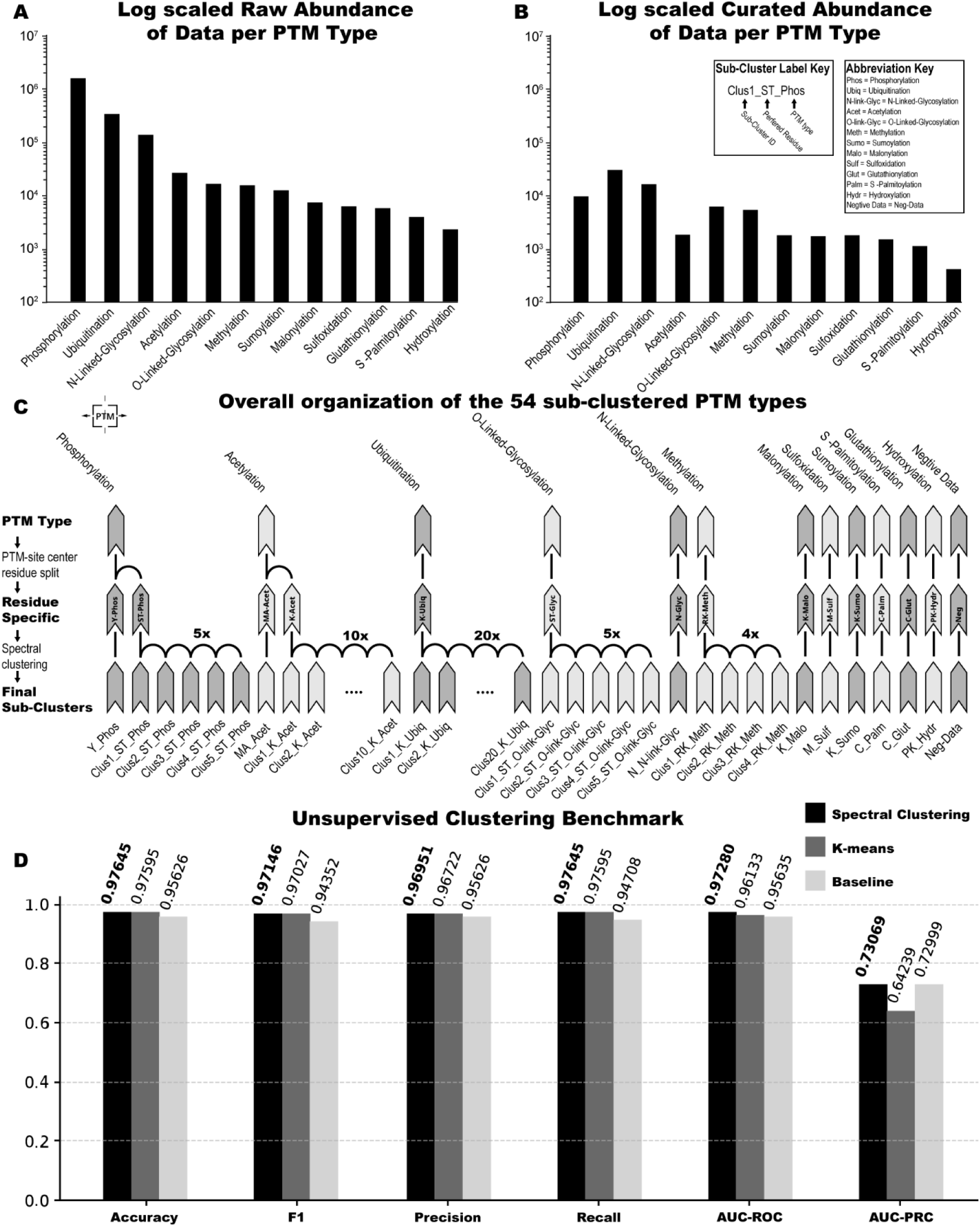
Overview of the CLASPP model’s data curation and stratification. (A) The log scale abundance of the data available for each PTM type pre-data curation for all species. (B) Log scale abundance of data available for each PTM type post-data curation for just human data. (C) The data hierarchy for each PTM type shows the organization of sub-clusters. Each sub-cluster is uniformly sampled to generate the final balanced dataset used for model training. (D) The choice of clustering algorithm consistently impacts the global macro PTM prediction performance. For this evaluation, a separate binary classification model was trained for each PTM sub-cluster using only human data. Performance was assessed using a 15% holdout validation dataset, enabling a fair comparison of the impact of different clustering methods on PTM prediction performance.

We next assessed the impact of different clustering methods on model performance. This includes k-means and spectral clustering applied to the peptide sequences of each PTM type as the input features. The number of sub-clusters is systematically varied (5, 10, 15, and 20) for the different PTM types with sufficient data abundance to allow for additional sub-clustering (S2 Fig). Spectral clustering displayed improvements in performance (F1=0.97146 and AUC_PRC=0.73069) compared to k-means (F1=0.97027 AUC and PRC=0.64239) and baseline conditions (F1=0.94352 and AUC_PRC=0.72999) (Fig 2c).

To evaluate the impact of clustering methods on model performance, we trained single binary classification models for each PTM type and the associated sub-clusters. This approach demonstrated that certain PTM types such as ST_Phosphorylation, K_Ubiquitination, K_Acetylation, and ST_O-Linked-Glycosylation are improved by sub-clustering (with downstream data stratification) (S2 Fig). The prediction performance of Y_Phosphorylation and N_N-Linked-Glycosylation is not impacted by sub-clustering. The remaining PTM types did not have sufficient positive data to support additional sub-clustering for this curation step. Other clustering quality metrics, such as mean silhouette coefficient [65] and the mean within cluster distances [66] were used to further evaluate the number of clusters per PTM type and the impact of different clustering approaches on model performance. These evaluations revealed that model performance was agnostic to clustering parameters for most PTM types, with the exception of Y_Phosphorylation, ST_O-Link-Glycosylation, and N_N-Linked-Glycosylation (S2c Fig.). These results demonstrate that implementing PTM-specific hierarchical sampling is an effective strategy to address data imbalance. This targeted sampling approach not only enhances the representation of each PTM type, but also contributes to more reliable and robust performance across diverse PTM types.

To assess whether the undersampling preserved the diversity of each sub-cluster, we examined both sequence-level and embedding-level representational spaces across all PTM types (S2f Fig.). For the sequence-based analysis, violin plots were constructed for each sub-cluster, with each half of the violin representing the distribution of pairwise average Hamming distances for each sequence among the full and undersampled populations of 21-mer input sequences. Each sequence in their respective populations was compared to every sequence in the full populations as a background. Near-identical distributions between the two halves were observed across sub-clusters for all PTM types. This indicates that undersampling did not introduce sequence-level distributional bias. Additionally, UMAP projections of the post-contrastive learning sequence embeddings were generated for each sub-cluster, with points colored by whether they belonged to the full or undersampled population. The embedding-space UMAPs similarly showed substantial overlaps between the two populations, with undersampled points distributed throughout the full-population manifold rather than confined to any particular region. Together, these analyses demonstrate that the undersampling strategy faithfully captures the sequence diversity and learned representational structure of each sub-cluster except for Y-Phosphorylation_nc0_tot1 and ST-Phosphorylation_nc0_tot5. The two problematic subclusters also had the largest reduction from full to subsample which is likely causing a large shift in UMAP representation and sequence identity distribution.

### Enhancing CLASPP with Contrastive Learning

We next integrated contrastive learning with the unsupervised PTM sub-clustering to learn nuanced features of individual PTM types. We introduced an additional contrastive learning stage between the traditional masked language modeling as a pre-training step and the final finetuning phase within CLASPP (S1 Fig). The contrastive learning stage of training is only performed on the ESM-2-150m encoder which is later used for downstream finetuning. The effects of two loss functions, Triplet Loss and Supervised Contrastive Loss, were systematically compared on downstream PTM predictions [64, 67]. We explored various finetuning regimes, specifically evaluating the outcomes when the ESM-2-150m encoder was either frozen or left trainable during the downstream finetuning. Our results demonstrate that the Supervised Contrastive Loss consistently outperformed both Triplet Loss and the baseline approach (Fig. 3a). Notably, maintaining the encoder in a frozen state during finetuning led to a slightly improved performance whenever contrastive learning was incorporated (Fig. 3a). These results highlight that supervised contrastive learning and encoder freezing during finetuning are efficient strategies to model multi-PTM events.

**Fig 3.**
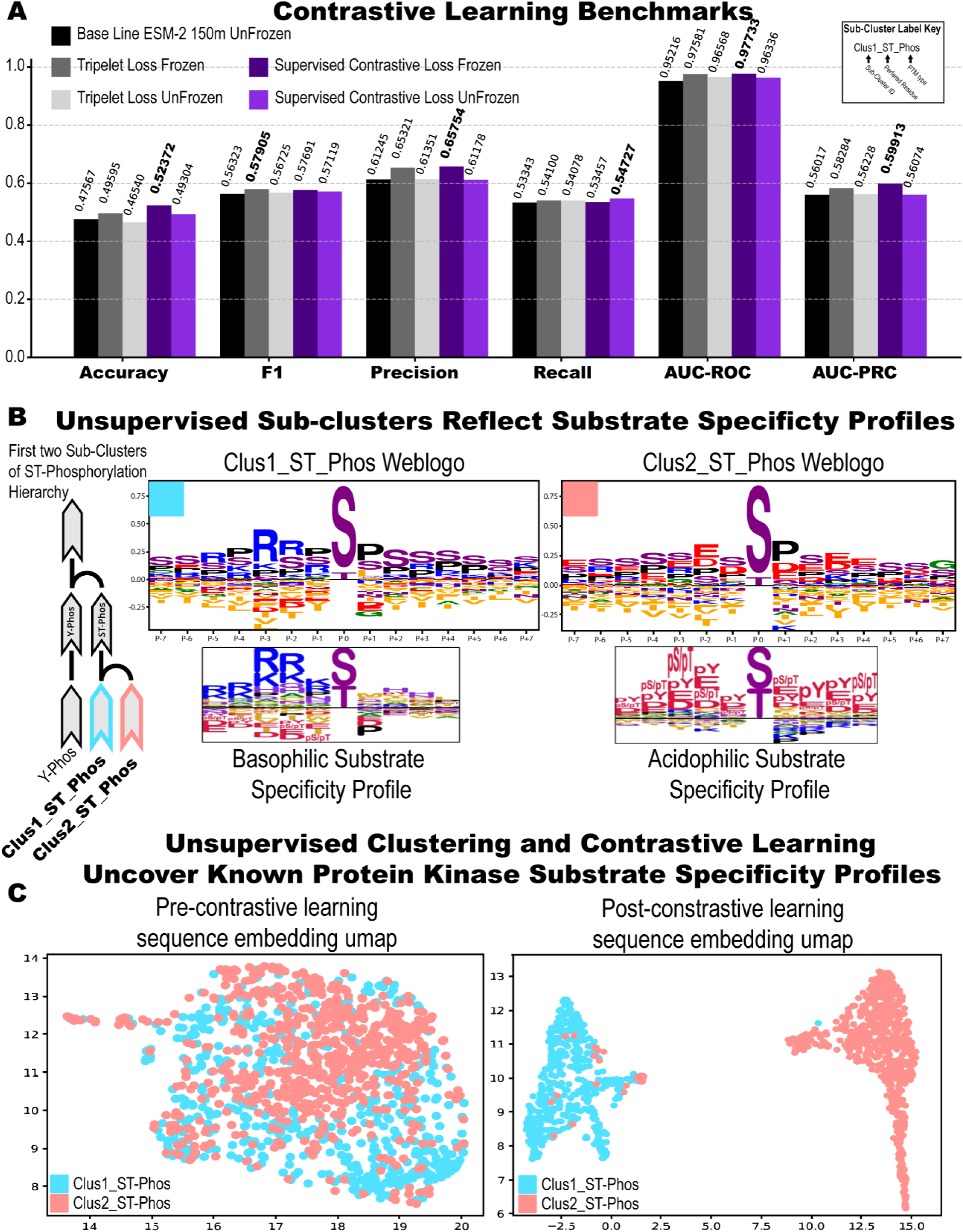
Investigation and explainability of the contrastive learning of the CLASPP model. (A) Comparison of the downstream global macro performance of the different contrastive learning scenarios. This benchmark only included human-associated PTM data and was measured with a 15% holdout validation dataset. (B and C) To investigate what the unsupervised clustering adds to the curation the first two sub-clusters found in the ST-Phosphorylation training dataset were enriched for kinase associations using Phosformer-ST [51]. In brief, the top 20 kinases were ranked by substrate abundance. The 21-mer peptide substrates of the top 20 kinases were pooled for each of the two sub-clusters to generate kinase-enriched dataset. (B) Weblogs of both the enriched sub-clustered datasets were compared with known protein substrate specificity profiles [55]. (C) A UMAP [68] of ST_Phosphorylation peptide embeddings pre and post contrastive learning. The enriched sub-clustered datasets were plugged into different checkpoints of the CLASPP model (pre and post-supervised contrastive learning). The outputs of the last layer of the encoder (embeddings) for each checkpoint are projected into a 2D space using UMAP with base settings. Additional parameter sweeps for UMAP and additional PTM types are also visualized across pre- and post-contrastive learning conditions. Generally, all PTM types tested resulted in better separation post contrastive learning (S3 Fig).

To determine whether the unsupervised clustering captures biologically meaningful features from the data, we visualize the amino acid preference of the peptide sequence in different enriched sub-clusters by generating sequence logos (Fig 3b). We focus on ST_Phosphorylation peptides because recent studies have grouped these peptides based on substrate specificity profiles of serine-threonine kinases that modify the central S/T residues based on amino acid preferences flanking the ST_Phosphorylation sites [55]. The sub-clustered peptides fall into categories that align with serine-threonine kinase substrate specificity. Sub-cluster 1 favors basophilic kinases, preferring arginine and lysine at the N-terminus of the central S/T residue (P-3, P-2 positions; Fig 3b), while sub-cluster 2 reflects acidophilic kinases, preferring aspartic and glutamic acids on both sides of the central phosphorylatable (S/T) residue (P-2 and P+3 positions; Fig 3b). Both clusters also show preference for proline at the P+1 residue of the phosphorylatable site, as demonstrated by previous experimental studies [55]. Because these amino-acid preferences in the peptide substrate are known to confer specificity towards an upstream kinase [69], the unsupervised clustering likely captures biologically meaningful representations.

To visualize how contrastive learning enhances sub-cluster separation, we projected sequence embeddings from pre and post-contrastive learning models using UMAP [68]. After all sub-clusters are established, the two enriched sub-clusters overlapped substantially in the UMAP projected embedding space. After contrastive learning, the enriched sub-clusters are distinctly separated when colored by their specific enriched sub-cluster labels which correlate with the substrate preference of upstream kinase involved in phosphorylation (Fig 3c). These results highlight that CLASPP, equipped with supervised contrastive learning, not only improves PTM prediction but also learns previously known biologically meaningful relationships among PTM types such as the substrate specificity of the kinase (PTM transferase) associated with substrate phosphorylation.

### Performance evaluation and comparison to other models

We next evaluated the performance of CLASPP by benchmarking against other multi-PTM prediction models such as MusiteDeep [18], MIND-S [19], PTMGPT2 [20], MTPrompt-PTM [25], and DeepMVP [21]. A uniformly sampled holdout test dataset, curated from unsupervised sub-clusters, was used for these evaluations. Benchmarks were performed on PTM types with sufficient data availability for model training and evaluation. These include Y_Phosphorylation, ST_Phosphorylation, K_Acetylation, K_Ubiquitination, ST_O-Linked-Glycosylation, N_N-Linked-Glycosylation, K_Methylation, K_Maloylation, K_Sumoylation, C_S-Palmitoylation, C_Glutathionylation, K_Hydroxylation, and P_Hydroxylation (Fig 4a). Notably, only CLASPP was restricted from accessing the test set during training.

**Fig 4.**
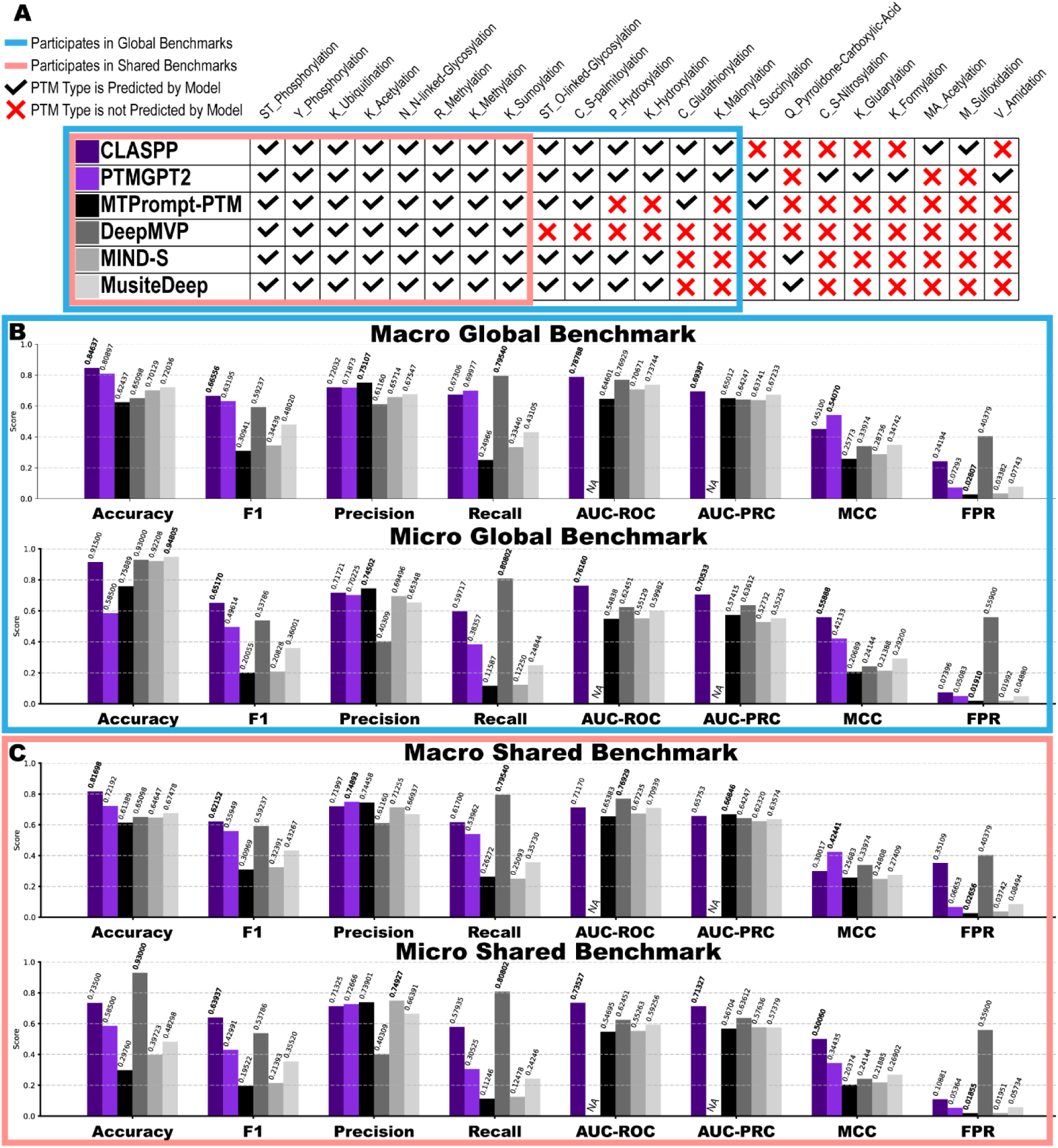
Global multi-PTM prediction benchmark. (A) Evaluation of models using global and shared benchmarks. Global benchmarks are based on all PTM types, while shared benchmarks are determined by PTM types commonly predicted by all models. Comparison of global and shared benchmarks for CLASPP, PTMGPT2 [20], MTPrompt-PTM [25], DeepMVP [21], MIND-S [19], and the MusiteDeep webtool [18]. The specific PTM types evaluated by each multi-PTM prediction model are indicated. (B) Presents the macro and micro global average PTM performance metrics. (C) Shows the macro and micro shared PTM type average performance metrics. (B and C) Benchmarks were conducted exclusively on human PTM data, using a 15% holdout test dataset. PTM types included in these benchmarks were selected based on having at least two models available for comparison and sufficient data to support CLASPP’s data curation process (see methods).

Across both the global (Fig 4b) and shared (Fig 4c) single label PTM benchmark performances, CLASPP consistently achieved state-of-the-art performance, though performance varied by PTM type. Overall, CLASPP globally outperformed other multi-PTM models specifically in Y_Phosphorylation (F1=0.74, and AUC_PRC=0.77), ST_Phosphorylation (F1=0.85 and AUC_PRC=0.94), and K_Hydroxylation (F1=0.61 and AUC_PRC=0.56). PTMGPT2 outperformed in R_Methylation (F1=0.92 and MCC=0.67), K_Methylation (F1=0.68 and MCC=0.68), ST_O-Linked-Glycosylation (F1=0.87 and MCC=0.77), C_S-Palmitoylation (F1=0.79 and MCC=0.65), P_Hydroxylation (F1=0.97 and MCC=0.69), K_Maloylation (F1=0.46 and MCC=0.5) and C_Glutathionylation (F1=0.94 and MCC=0.87). MTPrompt-PTM showed relatively better performance for N_N-Linked-Glycosylation (F1=0.96 and AUC_PRC=0.99) and K_Sumoylation (F1=0.6 and AUC_PRC=0.67). DeepMVP outperformed in K_Ubiquitination (F1=0.77 and AUC_PRC=0.86). An additional benchmark was performed using the MIND-S [19] test set to demonstrate performance on a traditional negatively sampled test set (See Methods). In short, the traditional negatively sampled test set favored models that have a low false positivity rate like MTPromptPTM, MIND-S, and MusiteDeep. All metric specific performance scores for all models and PTM types are shown in supplemental figure 4 in addition to confusion matrices for each model and benchmark(S4 Fig). Additionally, we benchmarked CLASPP and DeepMVP on the PTMAtlas test dataset as a form of extended validation (S5 Fig) [21]. Our evaluations indicate that PTMAtlas’s data tends to favor models with high sensitivity or high false positive rate, as evidenced by DeepMVP’s high recall and false positivity rate in Figure 4 and S5 Figure. Macro-based performance metrics, which normalize across PTM types, offer a less biased assessment than micro-based performance metrics that are influenced by data imbalance (Fig 4). Since the test data heavily features sub-clusters and PTM types related to lysine modification, such as K_Ubiquitination, K_Acetylation, and K_Methylation, micro global metrics show this bias.

Conversely, macro global metrics better reflect model performance by reducing the impact of overrepresented classes and offering a more balanced evaluation. Comparison of the micro and macro global performance of the different models on the dbPTM holdout test set shows that CLASPP generally performs better than, or as well as, other multi-PTM models in terms of accuracy, F1, precision, AUC_ROC, AUC_PRC, and FPR. Recall is the only metric that does not align with this pattern, as DeepMVP also exhibits a higher false-positive rate alongside its recall. We also compared model performance on predicting multiple PTM types on the same site by curating a multi-label dataset. In this evaluation, DeepMVP generally performed better than CLASPP in multi-label prediction (S4c Fig). In sum, these benchmarks provide an unbiased evaluation of the existing multi-PTM prediction models and offer guidance for future data curation and model evaluation studies.

### Evaluation of CLASPP in predicting PTM types across species and understudied kinases

Since the CLASPP model was only trained on human data, we next wanted to evaluate the performance of CLASPP on an independent out-of-distribution dataset of PTM types extracted from other model organisms (Fig 5a). Because of the non-uniform distribution of PTM types across species, we selected only those with at least 100 positive and 50 hard-negative data points for evaluation. A maximum of 500 randomly sampled positive and hard negative data points for each PTM type per species were used to maintain a balanced positive/negative ratio (Fig 5b). The PTM types Y_Phosphorylation and N_N-Linked-Glycosylation exhibit consistent performance across species (S6 Fig). Other PTM types show reduced performance based on accuracy, recall, and F1 metrics. Notably, precision, AUC_ROC, AUC_PRC, and FPR are consistent across species (Fig 5c) suggesting that positive predictions are likely to be correct predictions in other model organisms. Low recall and accuracy were observed in *Escherichia coli.* This suggests that the CLASPP model frequently misclassifies PTM types in this organism and that predictions need to be interpreted with caution in organisms that are evolutionarily divergent from the human (training) dataset.

**Fig 5.**
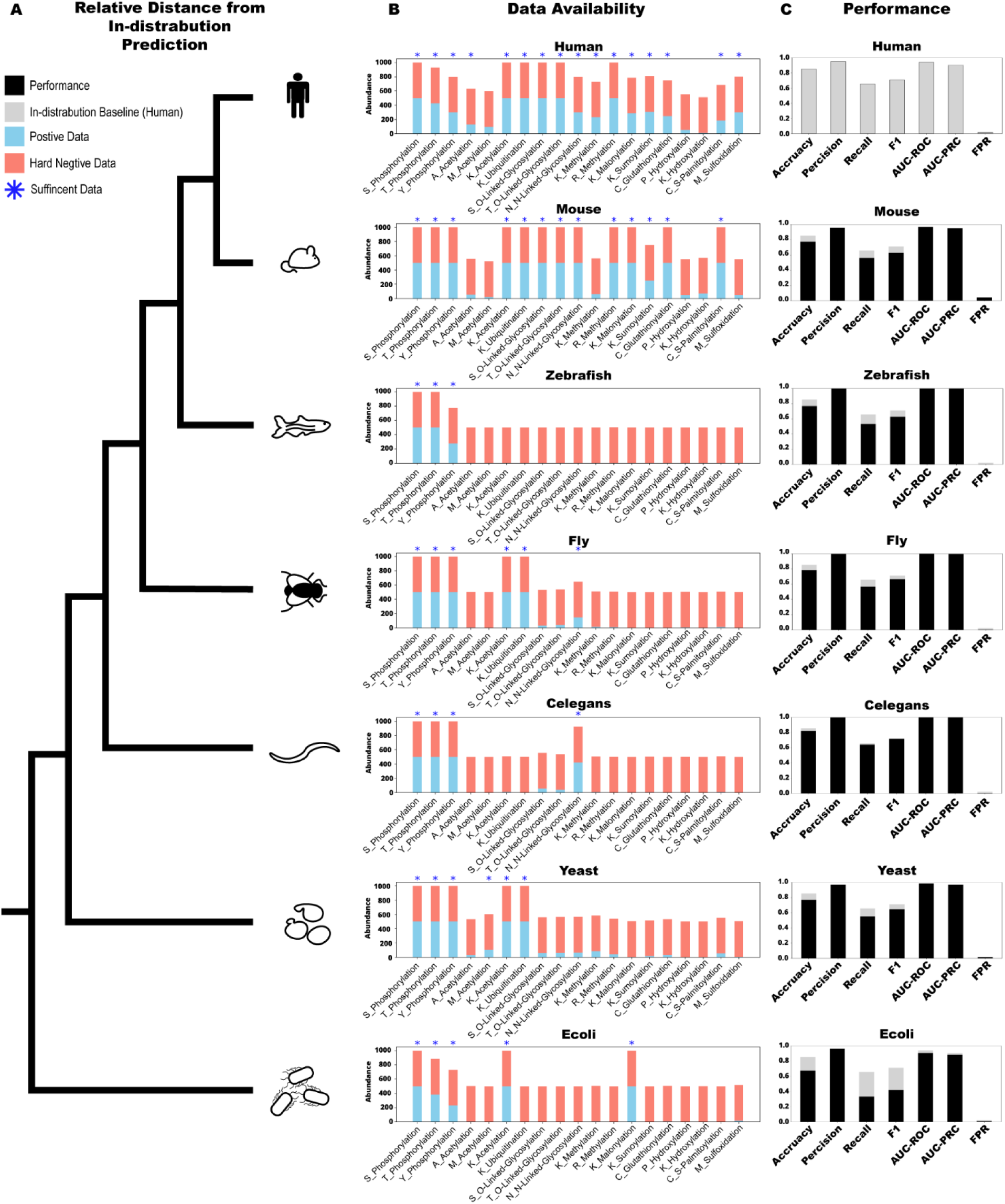
Comprehensive out-of-distribution benchmark of the CLASPP model. (A) Depicts relative evolutionary relationships of human and other model organisms used. Depicts the out-of-distribution evaluation of CLASPP model performance. Humans refer to *Homo sapiens*, Mouse refers to *Mus musculus*, Zebrafish refer to *Danio rerio*, Fly refers to *Drosophila melanogaster*, C. elegans refer to *Caenorhabditis elegans*, Yeast refer to *Saccharomyces cerevisiae*, and E. coli refer to *E. coli*. (B) The middle bar chart shows available data for each species-specific PTM type, marking those with at least 100 positive and 50 hard negative data points, with a blue star to indicate sufficient representation for evaluation. A cap of 500 randomly selected positive and hard negative data for each PTM type per species. (C) This out-of-distribution benchmark is limited to species-specific PTM types with sufficient data availability. The benchmarks present performance metrics based on the global macro average for species-specific PTM types with adequate representation, and compare them with the human baseline.

CLASPP also facilitates prediction of in-distribution (human) PTMs on understudied proteins such as DCLK3. The dark kinase DCLK3 is reported to exhibit very low levels of expression in various cell and tissue types and has recently been proposed to undergo ubiquitination-mediated degradation [63]. To illustrate the practical application of the CLASPP model, we compared its predictions of K-Ubiquitination sites with mass spectrometry validated sites not used in the training (Fig 6a). Of the five ubiquitination sites detected by mass spectrometry, only CLASPP predicted all of them correctly (Fig 6b). DeepMVP predicted four out of five reported K-Ubiquitination sites. PTMGPT2 and Mind-S predicted one site while MTPrompt-PTM and MusiteDeep predicted none. However, false-positive predictions are also high for CLASPP and DeepMVP compared to the other models.

**Fig 6.**
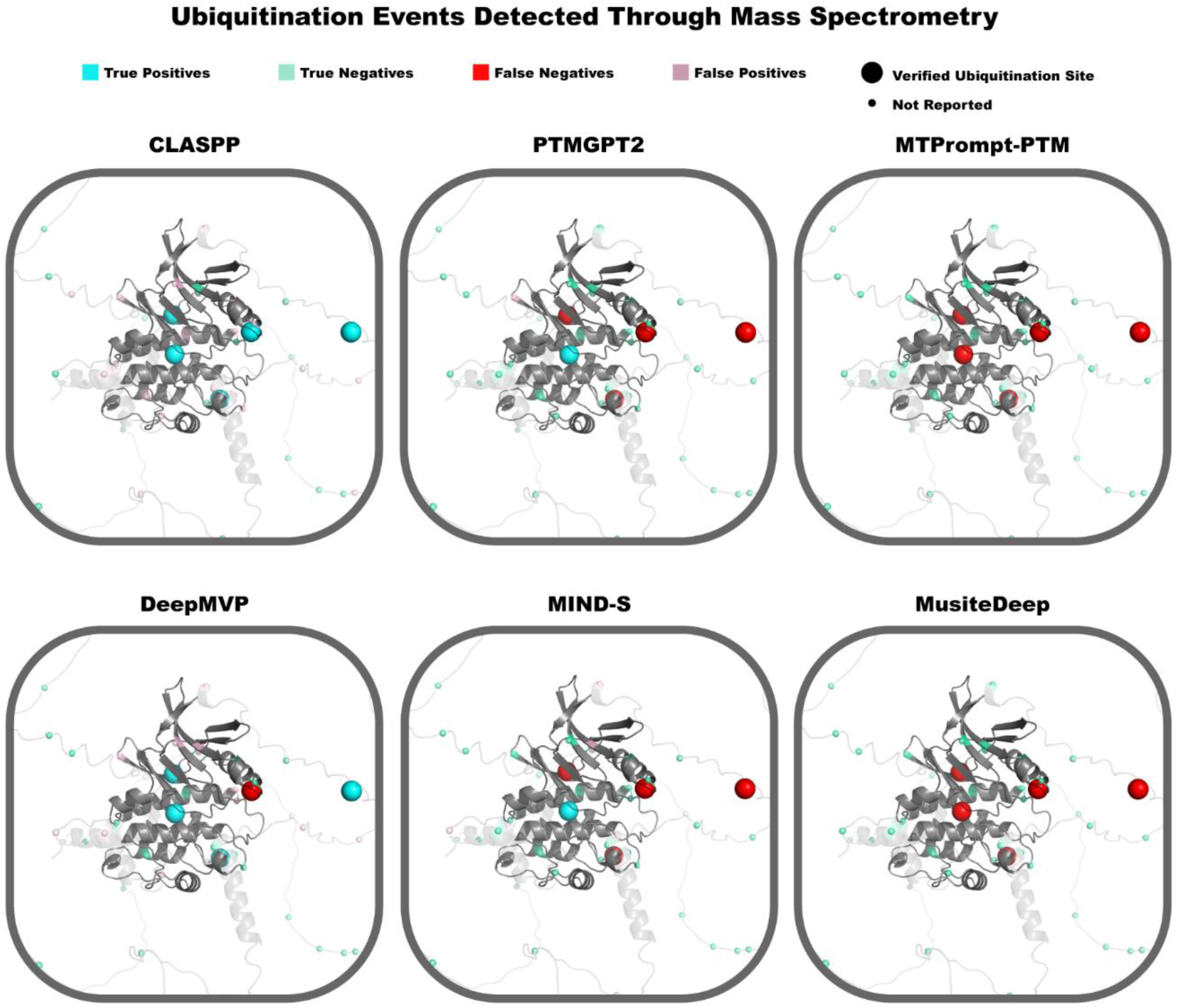
Overview of K-Ubiquitination site prediction in the understudied kinase DCLK3. Mapping of predicted K-Ubiquitination sites in DCLK3 as predicted by each model: CLASPP, PTMGPT2 [20], MTPrompt-PTM [25], DeepMVP [21], MIND-S [19], and the MusiteDeep webtool [18]. Lysine sites (shown as spheres) are mapped to the DCLK3 Alphafold 3D model (Q9C098) [70] using the Pymol software [71]. The protein kinase domain is shown in dark grey and the ground truth K-Ubiquitination sites were identified through Liquid Chromatography-Mass Spectrometry, described recently [63].

## Discussion

In this study, we benchmark existing multi-PTM prediction models and report on the development of the CLASPP model, which provides a unified framework for predicting multi-PTM events across species from sequence information alone [19–21, 25, 63].

CLASPP’s improved performance compared to existing models can be attributed to a few key factors, including the use of a protein language model that captures underlying biochemical and biophysical properties of proteins in an unsupervised manner by training on a large corpus of sequence data from diverse organisms [30]. PTMGPT2 employs a large language model, but it is not designed specifically for protein sequences [20]. Multi-PTM prediction models generally use all available data (after sequence identity filtering), which can lead to inflated performance on overrepresented PTM types due to overfitting [22]. Our data stratification strategies address this issue, and the unified model for predicting multiple PTM types offers a distinct alternative to previously proposed ensemble-based models [18–21, 25]. However, we note that we did not benchmark against recent models such as COMPASS-PTM because it appeared while our manuscript was under revision. Nevertheless, a key novelty of CLASPP compared to previous models is the deployment of multi-stage contrastive learning, enabling it to gradually learn PTM features [20, 25, 56]. In contrast, other PTM prediction models skip transfer learning and train all features simultaneously [19, 21]. These innovations make CLASPP more effective in multi-PTM prediction.

We demonstrate the application of CLASPP through two case studies. In our evaluation of CLASPP’s ability to predict PTM types across species, we demonstrate that it reliably predicts phosphorylation, acetylation, and N-linked glycosylation, but tends to miss true positives in out-of-distribution scenarios. Specifically, CLASPP displays a slight decrease in performance outside of Opisthokonta (Mouse, Zebrafish, Fly, Celegans, Yeast), but performance within metazoans and fungi is relatively stable. Stable performance is mostly observed for phosphorylation and acetylation across species, but not for other PTM types, presumably because the local sequence context is more conserved at phosphorylation and acetylation sites [72]. In addition, N-linked-glycosylation also shows comparable performance across species, consistent with previous studies [73]. Other PTM types displayed high precision but low recall, indicating few false positives yet low sensitivity. Consequently, while CLASPP predictions are reliable across species, CLASPP may miss true positives in out-of-distribution scenarios.

To demonstrate CLASPP’s effectiveness in predicting PTMs in understudied human proteins, we compared it with other multi-PTM prediction models for K_Ubiquitination sites on the human DCLK3 kinase, which were validated by mass spectrometry [63]. CLASPP correctly predicted all K_Ubiquitination sites but had a higher false positive rate than other models. This may be due to the lack of a strong sequence signal associated with these ubiquitination sites and/or the lack of true negative training data [9, 74]. Thus, future studies should focus on generating experimentally derived positive and negative training datasets to improve model performance.

CLASPP achieves state-of-the-art performance even with a 21-mer sequence window size, though other models like Mind-S [19] and AstraPTM [49] have shown performance increase with a larger context window size. However, given the current transformer-based architecture of the CLASPP model, increasing the context window size comes at a cost. While it may enhance the model’s understanding of long-range sequence relationships, it also increases the computational cost of training. Larger protein language models have demonstrated notable success in protein sequence classification (Sup. Fig. 1b). For instance, our experiments with ESM-3 and ESM-C show improved overall performance in PTM prediction. However, using models with more parameters as the baseline pre-trained protein language model significantly increases computational demands during additional training stages. To address these challenges, techniques such as layer freezing or LoRA [75] can be employed to reduce computational requirements. Furthermore, as demonstrated in our study, data stratification and curation strategies can also be viable approaches to improve model performance.

The optimal number of sub-clusters assigned to each PTM type is determined by both model performance and cluster-quality metrics. These two measures agree across PTM types with sufficient data to support additional sub-clustering. Among the three most abundant PTM types, the relative number of sub-clusters assigned to ST-phosphorylation (5), K-acetylation (10), and K-ubiquitination (20) is consistent with known differences in site-specific flanking patterns [76–78]. For phosphorylation, substrate recognition is driven largely by residues immediately surrounding the modified site [76, 79] which likely explains why this PTM requires the fewest sub-clusters among the three. This is consistent with the relatively constrained nature of protein kinase substrate specificity [55]. In contrast, acetylation appears to show weaker local site specificity around the modified lysine [77, 80]. Reported acetylation sites show greater diversity in surrounding amino acids, and their regulation is also influenced in part by cellular localization [81, 82]. Together, these features support an intermediate number of sub-clusters for acetylation. Ubiquitination, by contrast, is influenced by structural constraints that can extend beyond the immediate site, such as degrons [78, 83]. This suggests greater sequence diversity in sequences flanking ubiquitination sites compared to other PTM types, providing additional justification for why ubiquitination requires the largest number of sub-clusters to capture the prediction space effectively. For less abundant PTM types, limited data currently restricts the ability to sub-cluster them reliably. Within the current 21-mer context window, the optimal number of sub-clusters therefore appears to track with both regulatory complexity and sequence diversity around the PTM site.

Previous PTM prediction models commonly treated all unreported PTM sites as negative data [18, 19, 25], introducing a closed world assumption due to incomplete experimental coverage. In contrast, our curation strategy applies an additional constraint [84] by restricting negative samples based on sequence similarity to known PTM sites [32, 51]. Specifically, we enforce a minimum Hamming distance between candidate negative sites and all known PTM sites within a 21-mer sequence window. This constraint reduces false negatives arising from unannotated but sequence-similar modification motifs, thereby sharpening the decision boundary and improving robustness to label noise. We note that this sequence-similarity-based constraint is more appropriate for some PTM types than others. For example, phosphorylation is often governed by well-defined local sequence motifs, and thus benefits from this assumption [76]. In contrast, ubiquitination is influenced by degron context and distal structural features, which are not fully captured by local sequence similarity alone [78, 83]. As a result, the effectiveness of this constraint may vary across PTM types. Overall, while this approach does not uniformly model all PTM mechanisms, it represents a principled step toward mitigating the unrealistic assumption that all unreported PTM sites should be treated as true negatives.

As with many biological deep learning applications, PTM prediction is challenged by data abundance and class balance. Achieving robust performance requires careful representation of each PTM type in the training data and evaluation [25, 85]. As the volume of shared PTM data continues to grow, it is imperative to generate representative and curated datasets for training. PTMAtlas serves as a prime example: this dataset encompasses six distinct PTM types identified in human proteins [21]. Analysis of PTMAtlas and dbPTM reveals that 77% of the positive data points do overlap between the two datasets (Fig S5), highlighting substantial redundancy [9]. When benchmarking CLASPP and DeepMVP (which was trained on the PTMAtlas dataset), DeepMVP outperformed CLASPP. However, both models exhibited high false positive rates, which is also reflected in DeepMVP’s original dbPTM hold-out benchmark (Fig. 4). Combining both dbPTM and PTMAtlas into a unified dataset, combined with rigorous data balancing and curation techniques similar to those used in CLASPP’s curation, could substantially reduce the false positive rate and improve overall model performance. Likewise, expanding PTM data generation to include less represented modification types and sourcing data from a wider range of species beyond humans will be crucial. Additionally, sampling more easy negative data (see methods) may reduce false positive predictions but increase false-negative rate due to closed world assumption. Such efforts are expected to have the greatest impact on enhancing the quality and predictive power of multi-PTM models.

## Methods

To develop a unified model for PTM prediction, we first employed a protein language model as a baseline sequence classification architecture to predict 12 different PTMs types selected from the 70+ PTM types available in dbPTM. The selection of the 12 PTM types is based on the data abundance of the PTM type and past performance of previous multi-PTM prediction models [9, 18, 19, 21, 25]. All 12 PTM datasets were cleaned to the UniProt positional standard. After initial cleaning, the objective of the protein language model sequence classifier is to predict the PTM type from a 21-mer substrate sequence. Initially, we approached the task by training 12 separate PTM prediction models for each PTM type. This allowed us to establish performance baselines and refine our data curation strategies. Using the downstream performance from the individual models as a metric for quality, we employed a sequence identity threshold and evaluated the impact of unsupervised clustering. Based on the optimal clustering organization determined through benchmarking different scenarios of unsupervised clustering (see below), we uniformly sampled PTM 21-mer peptides to create a master list for training/testing/validation. Using the multiple sub-clusters as the 54 labels in the training/testing/validation sets, we trained a single unified PTM prediction model using contrastive learning. Finally, the unified PTM prediction model underwent a finetuning stage to predict the individual PTM sub-clusters.

### Data Sourcing

All data training/testing/validation is retrieved from dbPTM [9] and is verified using UniProt’s positional standard. Outdated UniProt accessions are recovered by using the UniParc database [11]. The training weights for the pre-trained ESM Protein Language Models were sourced from Huggingface [86] (ESM-2) and the ESM API [60] (ESM-3, ESM-C).

### Data Cleaning and Sequence Identity Threshold

To develop a robust training dataset for PTM prediction, careful curation of the 12 PTM types and their associated 21-mer modified sequences is required. First, we eliminated any sequences that did not exist or contained mismatched residue positions, ensuring all entries adhered to UniProt’s quality standards. Because much of the PTM data found in dbPTM are associated with outdated UniProt accessions, the UniProt REST API in combination with UniParc was used to accurately map these older accessions back to their correct sequences which maximized data recovery [11]. Additional data cleaning steps addressed inconsistencies in PTM site positions and updated position annotations with inconsistent numbering. Likewise, 21-mer peptide sequences that were non-human or lacked a canonical modified residue for each PTM type were removed from the dataset [17].

After initial cleaning, we employed representative greedy clustering methods to remove sequence identity redundancy [87]. This process entails filtering out highly similar 21-mer peptides based on sequence identity, using a Hamming distance cutoff of three. The greedy clustering was GPU-accelerated to efficiently run the program [88]. To ensure coverage of the sequence identity space, we randomly selected a unique representative from each cluster generated from the representative greedy clustering. For under-represented PTM types, we prioritized the random sampling of peptides from sequence identity clusters that contained multiple PTM types, thereby improving representation from less abundant modification types in the final dataset.

### Unsupervised Clustering

An additional unsupervised clustering curation step is added to address the non-uniform distribution of viable 21-mer peptides across PTM type. Without accounting for this disparity, PTM prediction models and evaluations will be biased towards over-represented PTMs in the database [22]. One way to solve the data imbalance issue is to split up the learning objective in an unsupervised manner, of certain PTM types, into more individual distributions. Unsupervised clustering achieves this goal and addresses the large range of prediction difficulty among all 12 PTM types. First the task is further broken up into 2-20 sub-clusters by using k-means clustering [58] and spectral clustering [59] on PTM types with sufficient data (enough to fully populate 2 separate sub-clusters). To plug into the scikit-learn pre-computed version of k-means and spectral clustering, individual affinity matrices for each PTM type were generated by creating the inverse of the Hamming distance matrix [65]. This implies that we use the 21-mer peptide itself as the input features. Ultra Scalable Spectral Clustering [89] is used to alleviate the computational constraints for just the ST-Phosphorylation dataset, due to the RAM usage of the scikit-learn version of spectral clustering. For only ST_Phosphorylation, we performed Ultra Scalable Spectral Clustering using 3D volumetric (Å^3) amino acid feature for the 21-mer peptides. Both scikit-learn and the Ultra Scalable Spectral Clustering only provide one sub-cluster label per 21-mer peptide and do not allow for data-leakage between sub-clusters. Finally, the best-performing number of clusters is determined across all PTMs by systematically benchmarking performance (S2 Fig). This subdivides the original 12 PTM type classes into 54 different PTM type sub-classes (Fig 2b).

### Uniform Sampling from the Sub-Clusters

Uniform sampling coupled with the unsupervised sub-clustering allows uniform data distribution across all PTM sub-clusters and allows for more complex PTM types to be accounted for. After the unsupervised clustering step, the total PTM examples were randomly sampled from each PTM sub-cluster, limiting the maximum number of positive examples to 2,000 per sub-cluster to reduce the over-representation of a small subset of abundant PTM types and sub-clusters. PTMs that had fewer than 2,000 examples would be fully utilized. All 53 positive and 1 negative examples are appended to a master list with a total of 97,989 PTM 21-mer peptides data points, and each peptide receives 54 binary labels. All 54 training sets are ensured to have even distribution across the training/validation/testing. The training/validation/testing stratification sampling was reserved for the last step in the curation to ensure no leakage across the sets.

Consequently, the training/validation/testing all have an even distribution of the 54 sub-clusters. The overall training/validation/testing split is 70%/15%/15%, respectively.

### Multi-Label Handling of The Multi-PTM Data

Prior to down-sampling of the 54 sub-clusters, a priority sampling is performed for each PTM sub-cluster, ensuring that multi-PTM event (peptides associated with more than one PTM type) data points comprise up to 20% of the 2000 total sub-sampled peptides for each of the 53 positive sub-clusters. This was done to reduce the oversaturation of multi-label data in our datasets. This 20% cap of multilabel peptides is applied to all PTM sub-clusters except for Malonylation and Sumoylation due to naturally high participation in multi-PTM events in dbPTM for humans [90].

### Negative Data Handling

We employed a negative data handling strategy described in our previous work [32]. In brief, we introduced three negative data categories (hard negative, medium negative, and easy negative) and employed prioritized negative data sampling to address data imbalance within each PTM type. The three negative categories were defined as follows: hard negatives are defined by cluster assignment. For example, for the class Clus1_ST_Phos, data points from the remaining 53 sub-clusters would be considered hard negative. Medium negative data corresponds to human PTM types found in dbPTM but not to the 12 PTM types of interest. Easy negative data points are randomly sampled 21-mer peptides in the full protein sequence for which there is no experimental evidence of a modification. Each easy negative data point is at least a Hamming distance of 7 away from any known positive PTM example in the dbPTM of any species. It should be noted that, most other PTM prediction models use only a non-filtered version of easy negative data [85]. Our approach is unique in that it considers hard and medium negatives and adds an additional filter to the easy negative sampling. Traditional negative sampling consists of non-reported residues that share a residue type. For example, if a protein had 10 lysine residues and 2 of them are reported ubiquitination sites, the remaining 8 lysine sites are considered negative data.

For the construction of the benchmark datasets used in figure 4 (hold-out test set) for every PTM type, we design the benchmarks to be residue specific to match the input constraints of the other models [18, 19, 25]. For the positive datapoints, this includes all sub-cluster specific positive data associated with each PTM type. For the negative datapoints we included any hard negative data that share the residue type. The rest of the negative data is derived from the medium and easy negative samples that share the residue type of interest. By constructing the PTM benchmarks this way, we ensure even representation from each sub-cluster.

Additionally, we developed datasets with varying positive/negative ratios. We also generated datasets with varying ratios of modified residues. The final multi-binary classification model would not converge if the negative ratio was too high per PTM sub-cluster. In response, we capped the negative ratio to 1/60 in the final model and prioritized negative data from the medium negative class over easy negative class for the single negative class. We also ensured that all data for the easy negative label had a uniform distribution across the residue types of the potential modified residue (the center residue of the 21mer peptide).

### Model design

After a series of benchmarks of different ESM-based architectures [30, 60, 61], our final architecture comprises a pretrained ESM-2-150m encoder followed by a two linear layer classification head, yielding 54 independent binary logits for multilabel PTM prediction (S1 Fig). Each output logit represents the model’s prediction on the different sub-clustered PTM types. 21-mer peptides are fed into the ESM encoder with full embedding (including <cls> and <eos> tokens) used as the input for the multi-label binary classification head. The model’s final training is done with a loss function that includes a sigmoid function that turns the final logits into probabilities. The ESM-2-150m encoder architecture weights were first downloaded from Huggingface, already pre-trained from the mask language modeling step [30], and were later put through a supervised contrastive learning task [64] with the 54 PTM sub-classifications as the dataset that was also used for the finetuning stage. Finally, a multi-binary classification finetuning step is performed, with all parameters in the ESM-2-150m encoder kept frozen to preserve knowledge from the multiple pre-training steps. Once training is complete, predictions for each sub-cluster class are aggregated using max pooling for each residue specific PTM type (e.g., S_Phos, T_Phos, Y_Phos, K_Ubiq, etc.). This process yields a final output of 19 positive labels and 1 negative label, for a total of 20 labels. Max pooling is used for easier comparisons in the benchmarking.

### Contrastive Learning

To evaluate how contrastive learning can incorporate proximal PTM class information into the protein sequence latent variable space, we used two distinct loss functions: Triplet Margin Loss [67] and Supervised Contrastive Loss [64]. Both approaches were systematically benchmarked against ESM-2-150m baseline.

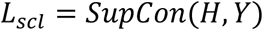

Let *L*_*scl*_ denote Supervised Contrastive Loss [64]. Let *H* denote the batched embeddings deriving from the output of the last layer of the ESM-150m encoder. Additionally, *H* includes sequence and special tokens. And let *Y* denote the batched sub-clusters labels. Baseline hyperparameter settings are used (S1a Fig.).

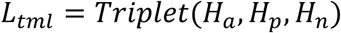

Let *L*_*tml*_ denote Triplet Margin Loss [67]. Let *H*_*a*_ denote the batched embeddings deriving from the output of the last layer of the ESM-150m encoder for the anchor sequences. Let *H*_*p*_ denote the batched embeddings deriving from the output of the last layer of the ESM-150m encoder for the positive pair sequences. Let *H*_*n*_ denote the batched embeddings deriving from the output of the last layer of the ESM-150m encoder for the negative pair sequences. Additionally, *H*_*a*_, *H*_*p*_, *H*_*n*_ includes sequence and special tokens. Baseline hyperparameter settings are used (S1b Fig.).

The data used with the Triplet Loss is prepared by iterating through all positive data points for each of the 54 PTM sub-classes. Positive and negative pairs are randomly sampled with a set of constraints. 1) The first constraint for each Triplet Loss sample is checking if the anchor-positive pair and the anchor-negative pair have been sampled before to prevent redundancy. 2) The next constraint is that each anchor point can only have a maximum of 150 positive and negative pairs to reduce the exponential scaling of all possible permutations. 3) The final constraint consists of the positive pairs that are sampled from the same sub-cluster, where the negative pairs are sampled from different PTM types altogether. This means data that falls in the same PTM type, but different sub-clusters is not used as a negative pair. We also altered the data-loader, loss function, and batch sizes (3x) to accommodate the Triplet Loss contrastive learning training objective.

As for the Supervised Contrastive Loss, the data structures, data-loader, or batch-sizes were not altered from the finetuning stage of training. Only the loss function of the model is altered to accommodate the Supervised Contrastive Loss learning objective.

For both contrastive learning tasks, the checkpoints of interest are measured by their downstream performance in the finetuning task of PTM classification. During the contrastive learning stage, overfitting was assessed solely through training and validation loss. Validation loss served as the preliminary criterion for checkpoint selection, and the subsequent checkpoints were verified by training a classifier on each checkpoint to determine the best-performing model.

### Finetuning

The final finetuning training step involved a multi-label sequence classification task across 54 distinct PTM sub-clusters. Each input was a 21-mer protein peptide sequence, and the output is a probabilistic prediction for each of the 54 PTM sub-clusters. Binary Cross-Entropy was employed as the loss function to optimize the model for this multi-label PTM prediction task. Finetuning is used to determine the downstream performance for the curation, model, and training design choices. We conduct an extensive evaluation of various possible method combinations to identify the best-performing approach. To convert continuous prediction scores into binary outcomes, we apply a threshold of 0.5 for the metrics of accuracy, F1, precision, recall, MCC, and FPR. All metrics used were calculated using scikit-learn [65]. When evaluating the unsupervised clustering section (Fig. 2 and SFig. 2), we used a single binary classifier for each PTM type or sub-cluster to better isolate variables.

### Hyperparameters and Hardware

For the contrastive learning stage, we employ a learning rate of 2e-6 with a decay of 0.001, which was found to be stable. The training was conducted with a batch size of 124 with a gradient accumulation of 4. Training for 1000 epochs was sufficient for convergence. All training and validation were performed on four Nvidia RTX 6000 Ada(s) in parallel [91]. Under these conditions, the training for the contrastive learning stage was less than 72 hours.

For the finetuning stage of the final model, we employ a learning rate of 2e-6 with a decay of 0.05, which was found to be stable. The training was conducted with a batch size of 124 with a gradient accumulation of 4. Training for 500 epochs was sufficient for convergence. All training and validation were performed on a single Nvidia RTX 6000 Ada. Under these conditions, the training for the finetuning stage was less than 24 hours.

## Data Availability and Code

The Github for the CLASPP model is available here (https://github.com/gravelCompBio/Claspp_forward). The Huggingface is available here (https://huggingface.co/esbglab/Claspp_forward). Training datasets for both finetuning and contrastive learning are available here in this Zenodo repo(https://zenodo.org/records/17674057) and each benchmark is mapped to what figure it was used for. The Github of the Data Curation is available here (https://github.com/gravelCompBio/Claspp_data_cur).

## Author Contributions

N.G., Z.Z., and N.K. came up with the **Conceptualization** of this project. N.G. oversaw the **Data Curation** for the project. N.G., Z.Z., N.K., and R.F. designed and implemented the **Formal Analysis** for this project. N.K. oversaw the **Funding Acquisition.** N.G., R.F., Z.Z., S.S., and N.K all contributed to **Methodology.** N.G., R.F., A.D., and N.K. contributed to **Resources** management for this project. N.G., R.F., Z.Z., S.S., and A.D. developed all associated **Software.** N.G., N.K., R.F., Z.Z., S.S., and A.D. all contributed to the **Writing Preparation and Editing** along with the **Visualization.**

## Funding Acknowledgement

Support for NK from the National Science Foundation (DBI-2400220; NSF BioF: GREAT) and the National Institutes of Health (R35-GM139656) is acknowledged. The funders did not play any role in the study design, data collection and analysis, decision to publish, or preparation of the manuscript.

## Supporting Information

**S1 Fig. Model training and architecture overview**

(A) Overview of the supervised contrastive loss training scheme. (B) Overview of the triplet loss training scheme. (C) Overview of the fine tuning training scheme. (D) The first contrastive learning step was performed on the encoder. (E) Then the classification head was finetuned. (F) Overview of the ESM-2’s encoder’s multiheaded attention [30]. (G) Overview of the attention mechanism. (H) Overview of the classification head used. (I) Investigation of the different encoders and classification head architectures. This uses macro performance across the holdout validation set. This benchmark was trained without a contrastive learning step and only a finetuning step. Identical training regimens were applied to each architecture. 2LinearClassifer refers to the architecture you see in S1e Fig and 3X8Transfomer refers to an architecture where you swap the first linear layer with a multiheaded attention layer that is 8 wide and 3 deep [49]. We chose ESM-2-150m due to the comparably superior performance and due to computational constraints of ESM-2-150m being the largest model we can feasibly use for contrastive learning. The same hyper-parameters were used for all architectures benchmarked (originally optimized with ESM-2-150m) and could be potentially optimized for ESM-C-300m, ESM-C-600m, and ESM-3-open individually [60, 61].

**S2 Fig. Comprehensive unsupervised clustering benchmark and by PTM type and data curation overview**

Systematic benchmark of the differing number of clusters and clustering method. (A) Left column is reserved for baseline performance without unsupervised clustering. (B) Middle column is reserved for the supervised clustering benchmarks. (C) Right column is reserved for clustering quality metrics which includes mean silhouette coefficient (silhouette) and mean within cluster distance (within cluster distance). Each PTM type was trained and validated using a unique single binary classification model for each sub-cluster or PTM type. The performances of each sub-cluster and their individual model were pooled together using global macro performance to represent the assigned number of sub-clusters. (D) Overview of the data curation consisting of downloading the data from dbPTM, cleaning according to UniProt standards [11], and filtering for human and residues most associated with the PTM type of interest. Followed by a sequence identity threshold to ensure there is an even distribution across sequence identity space. Then a spectral clustering applied to each PTM type with benchmarked proof that it increases performance and allows for a more diverse sequence identity distribution per PTM type [55, 73]. Finally, a uniform sampling and training/testing/validation split is performed [22]. (E) A PTM specific abundance table for various stages of the data curation pipeline between steps that filter or down sample the known PTM sites. (F) Violin plots and UMAP plots demonstrate how the undersampling preserved the diversity of each sub-cluster. The violin plot measures each sequence’s average hamming distance away from the full population and shows full and undersample distribution. The UMAP shows the post-contrastive embeddings of each data point.

**S3 Fig. Systematic visualization of UMAP projections of all participating PTM types pre/post-contrastive learning across different parameters of UMAP**

UMAP representations of pre- and post-supervised contrastive learning training schemes. These tests are divided into 7 different PTM-centric populations including global, phosphorylation, ubiquitination, acetylation, glycosylation, methylation, and others. This test was replicated over different input parameters that impact UMAP projections like random seed, n_neighbors, and min_dist. This data was pulled from the test dataset (was not used in training).

**S4 Fig. Comprehensive benchmark performance from all PTM types**

All PTM types that have at least 2 models associated with the PTM type are tested. Prediction generated from CLASPP, PTMGPT2 [20], MTPrompt-PTM [25], DeepMVP [21], MIND-S [19], and the MusiteDeep webtool [18] for all PTM-types. For the different PTM types, we measured the multi-label performance by including all positive testing data and negative testing data that belongs to the same residue type for the hard, medium and easy examples. F1 and AUC_PRC are the primary metrics to determine overall performance for discrete (threshold of 0.5) and probabilistic modeling respectively. S_Phosphorylation T_Phosphorylation are tested separately, and the mean of the two scores are shown. S_O-Linked-Glycosylation and T_O-Linked-Glycosylation are tested separately and the mean of the two scores are shown. This pooling (mean) was done for ST_Phosphorylation and ST_O-Linked-Glycosylation to account for the residue specific way of modeling multi-PTM prediction. (A) Benchmark of the hold-out data from the CLASPP tests. (B) Benchmark of the hold-out data from the MIND-S tests. (C) Overview of multi-label performance benchmark. Comparison of shared PTM types that participate in multi-label events for CLASPP, PTMGPT2 [20], MTPrompt-PTM [25], DeepMVP [21], MIND-S [19], and the MusiteDeep webtool [18]. The specific PTM types evaluated by each multi-PTM prediction model are indicated. The bar chart shows available data that shares the presented labels and denotes which multi-label benchmark has enough data to be significant with a blue star. Any benchmark that had 30 or more positive data points was considered significant. Accuracy score, Jaccard score, mean absolute score was imported from sk-learn [65], and the metric accuracy only considers a prediction correct if it matches every single ground truth label. (D) Global and shared benchmarks of the hold-out data from the CLASPP and MIND-S test-set. (E) Confusion matrices generated for all individual models and test set.

**S5 Fig. Comparison of PTMAtlas and dbPTM**

(A) Probability of PTMAtlas overlapping with dbPTM of positive 21-mer sequences per PTM type [9, 21]. (B) Measure of sequence diversity per PTM type for PTMAtlas and dbPTM. The metric of sequence diversity was measured by the log scaled number of clusters from a representative greedy clustering of the positive 21-mer sequences. Different thresholds were systematically tested. (C) Benchmarks of DeepMVP [21] and CLASPP ran on PTMAtlas’s testing set. Binary-label benchmarks refer to using the exact test data supplied by PTMAtlas. Multi-label benchmarks are the altered test sets where the positive data points from adjacent PTM types are added to the negative class of interest so long as the 21-mer input does not exist in the positive class of interest and they share the central modified residue type.

**S6 Fig. Comprehensive out-of-distribution benchmark.**

(A) Demonstrates relative evolutionary distance and this metric is a way to measure the degree of out-of-distribution for a given species-specific benchmark. Human refers to *H. sapiens*, Mouse refers to *M. musculus*, Zebrafish refers to *D. rerio*, Fly refers to *D. melanogaster*, Celegans refers to *C, elegans*, Yeast refers to *S. cerevisiae*, and Ecoli refers to *E. coli*. (B) This out-of-distribution benchmark is for the metric accuracy. (C) This out-of-distribution benchmark is for the metric F1. (D) This out-of-distribution benchmark is for the metric precision. (E) This out-of-distribution benchmark is for the metric recall. (F) This out-of-distribution benchmark is for the metric AUC-ROC. (G) This out-of-distribution benchmark is for the metric AUC-PRC. (H) This out-of-distribution benchmark is for the metric FPR. (C-H) The benchmark only contributes to the Fig 5c if a blue star is present representing sufficient data.

## Notes

https://zenodo.org/records/17674057

https://github.com/gravelCompBio/Claspp_forward

https://github.com/gravelCompBio/Claspp_data_cur

### Competing Interest Statement

The authors have declared no competing interest.

