## Supplementary figures and images for "CLASPP: A unified model for predicting post-translational modifications"

### S1Fig.png

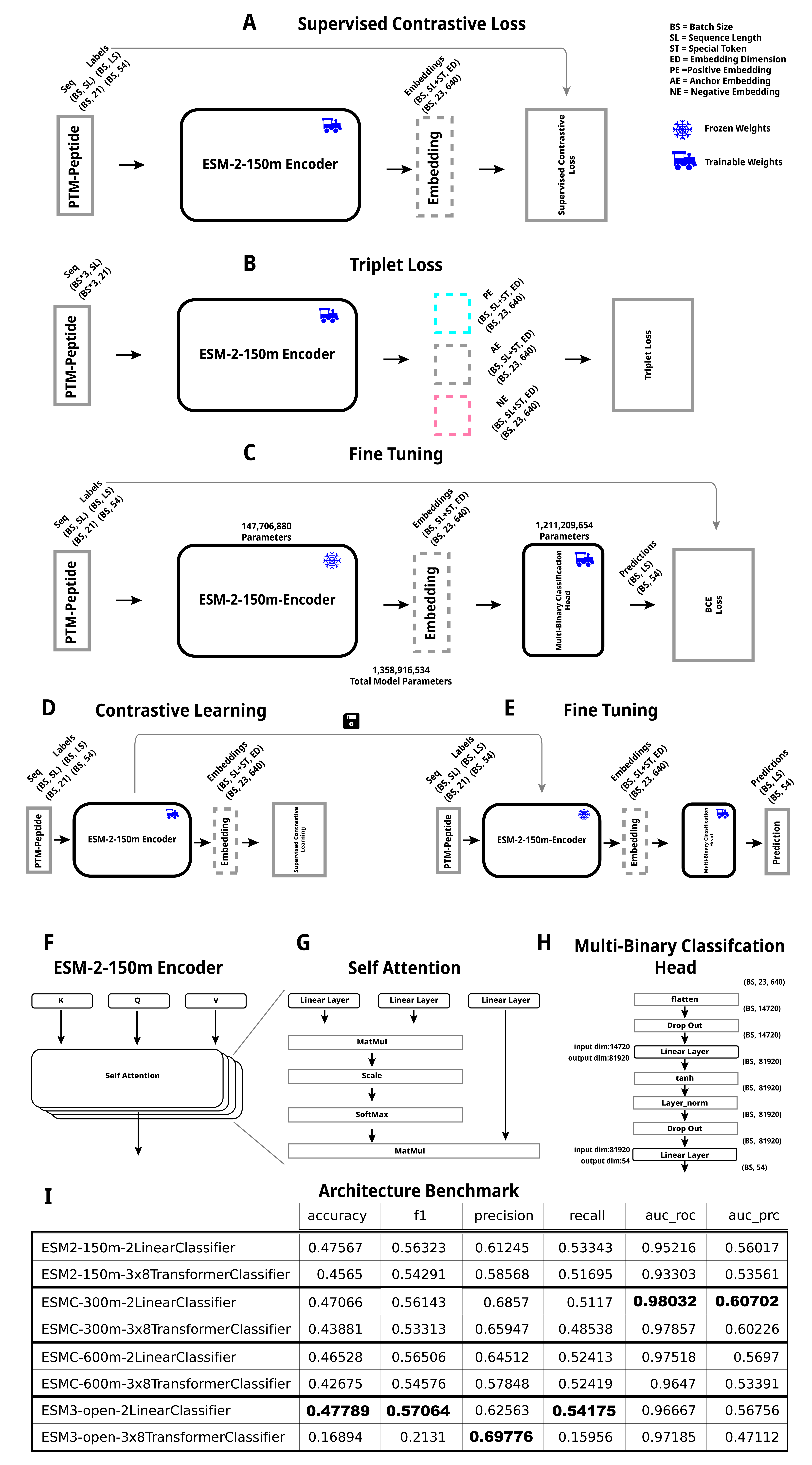

### S2Fig.png

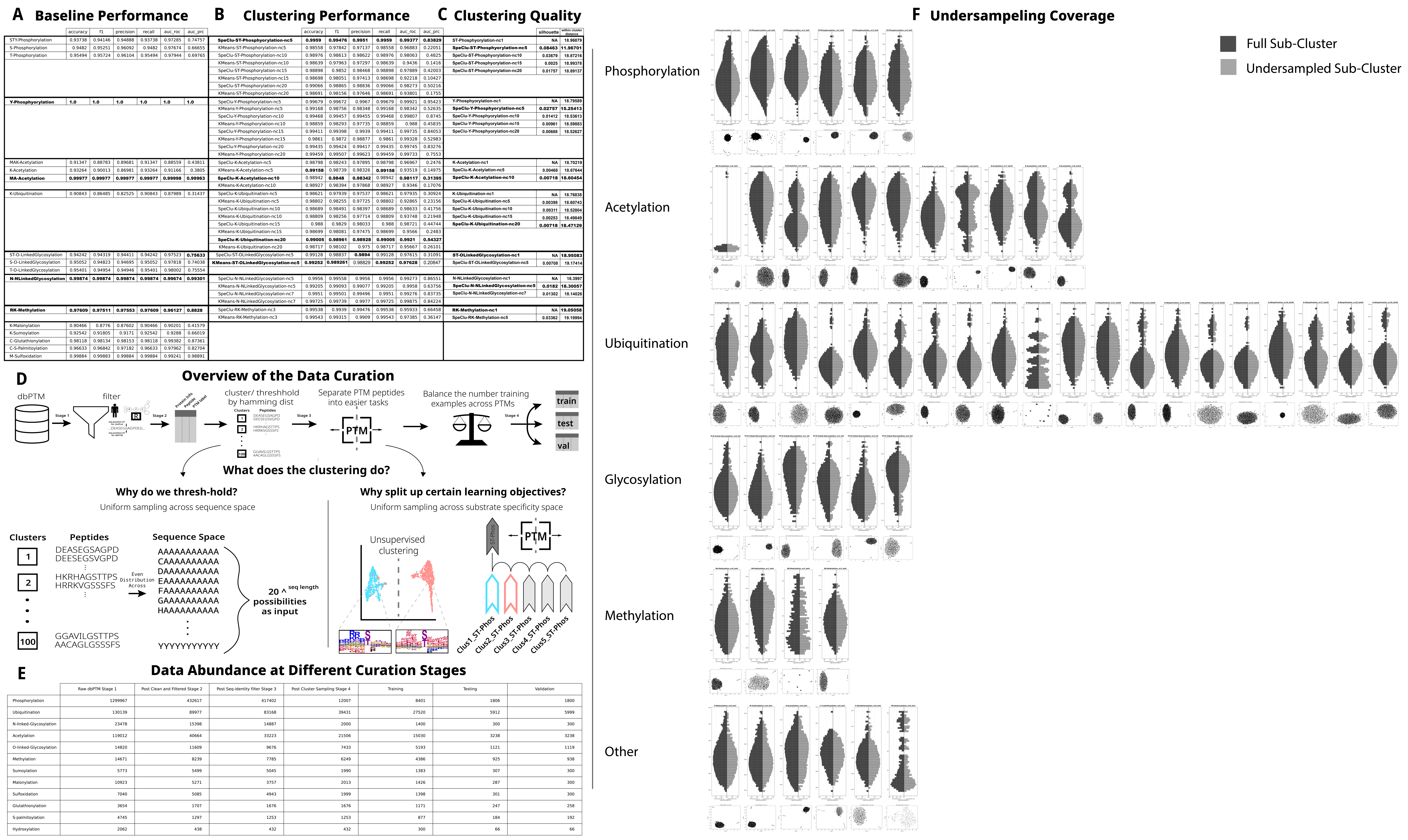

### S3Fig.png

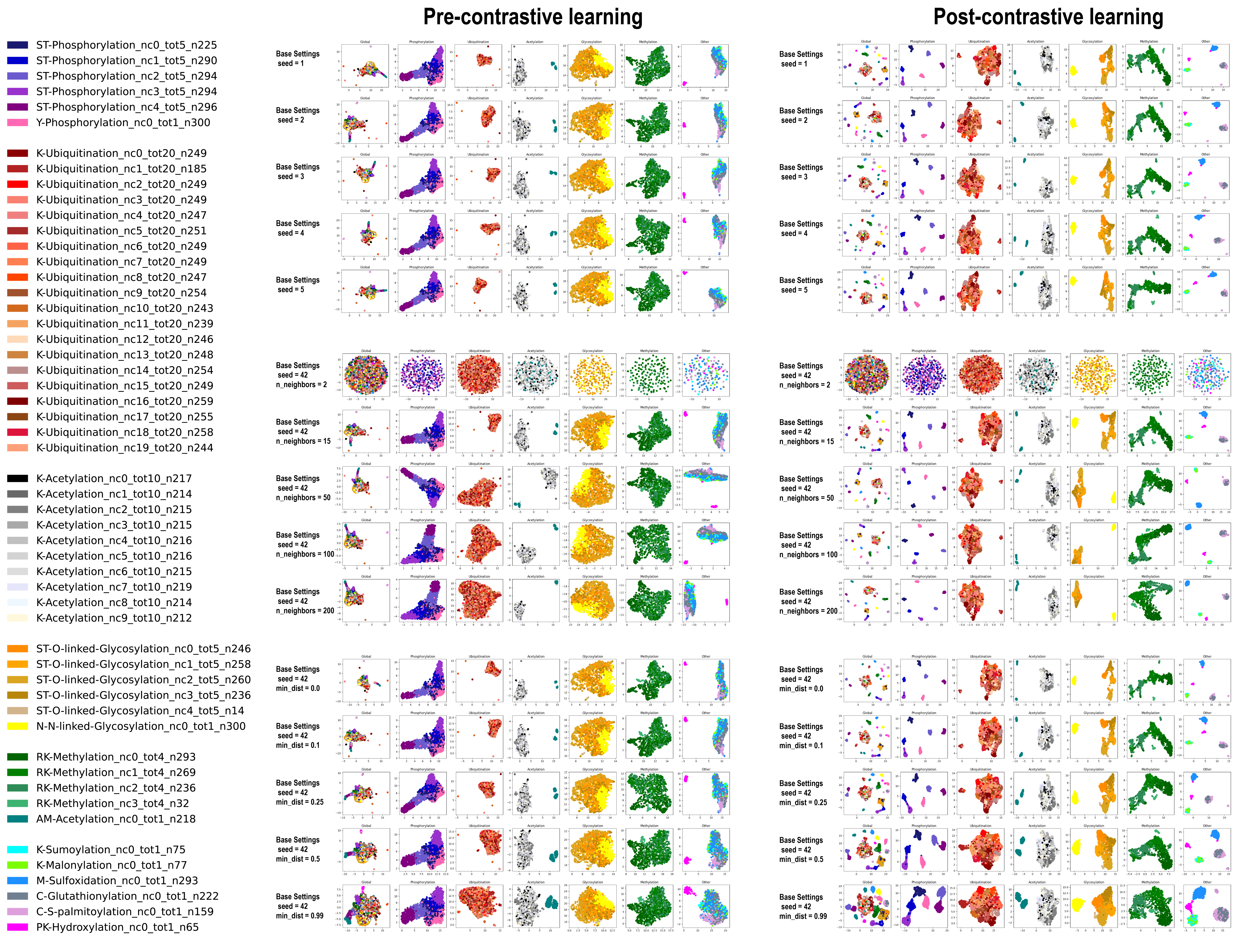

### S4Fig.png

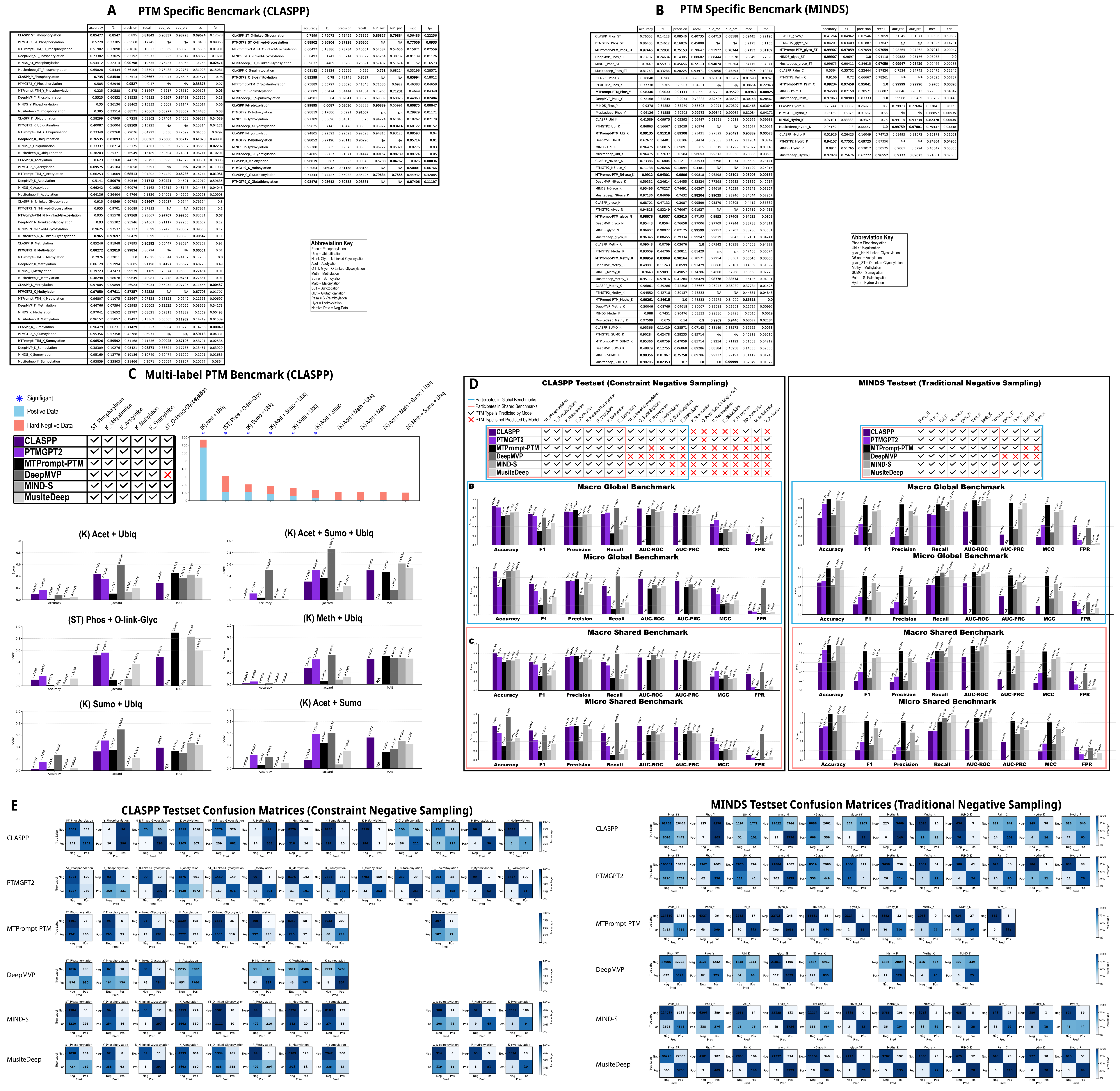

### S5Fig.png

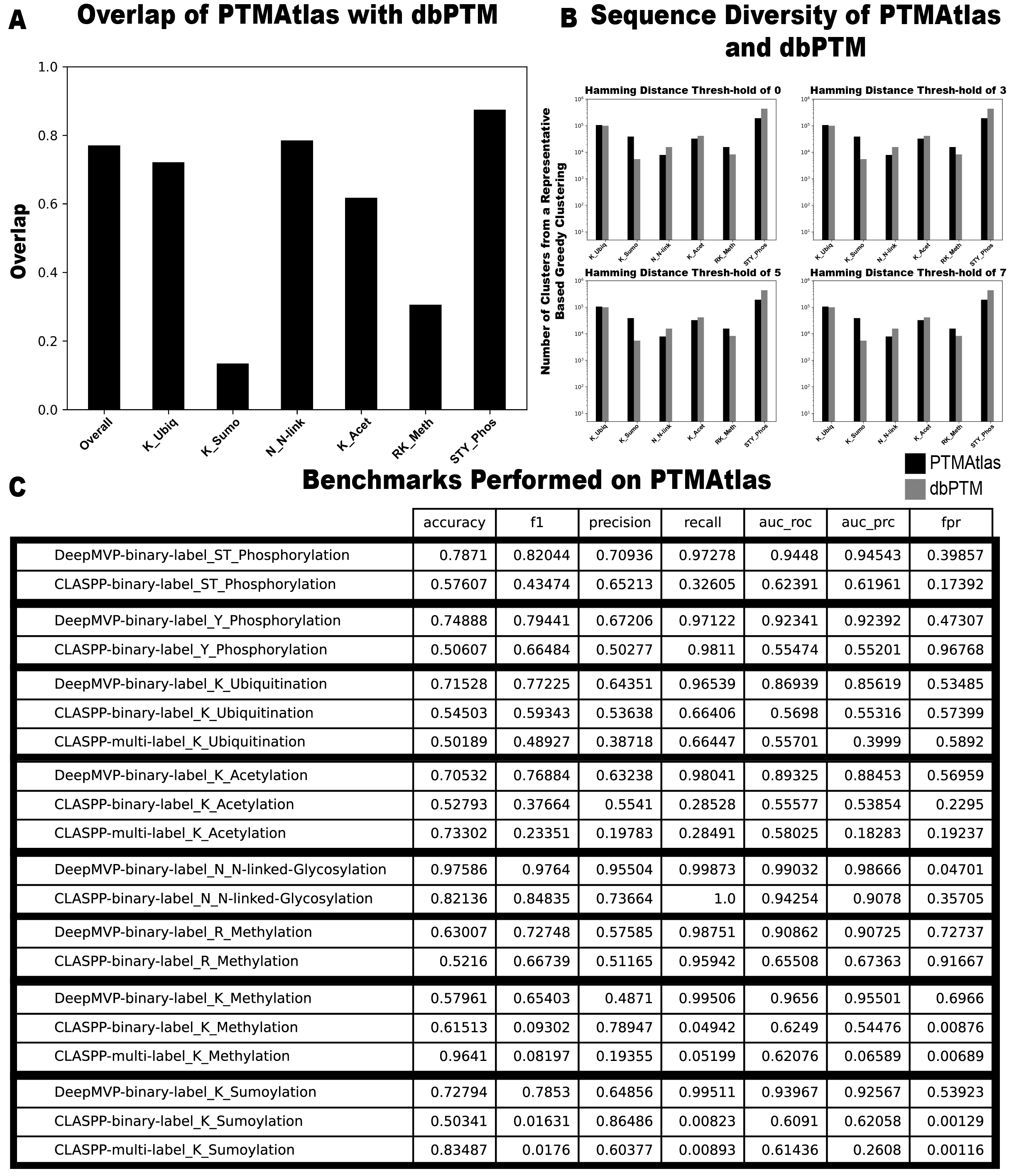

### S6Fig.png

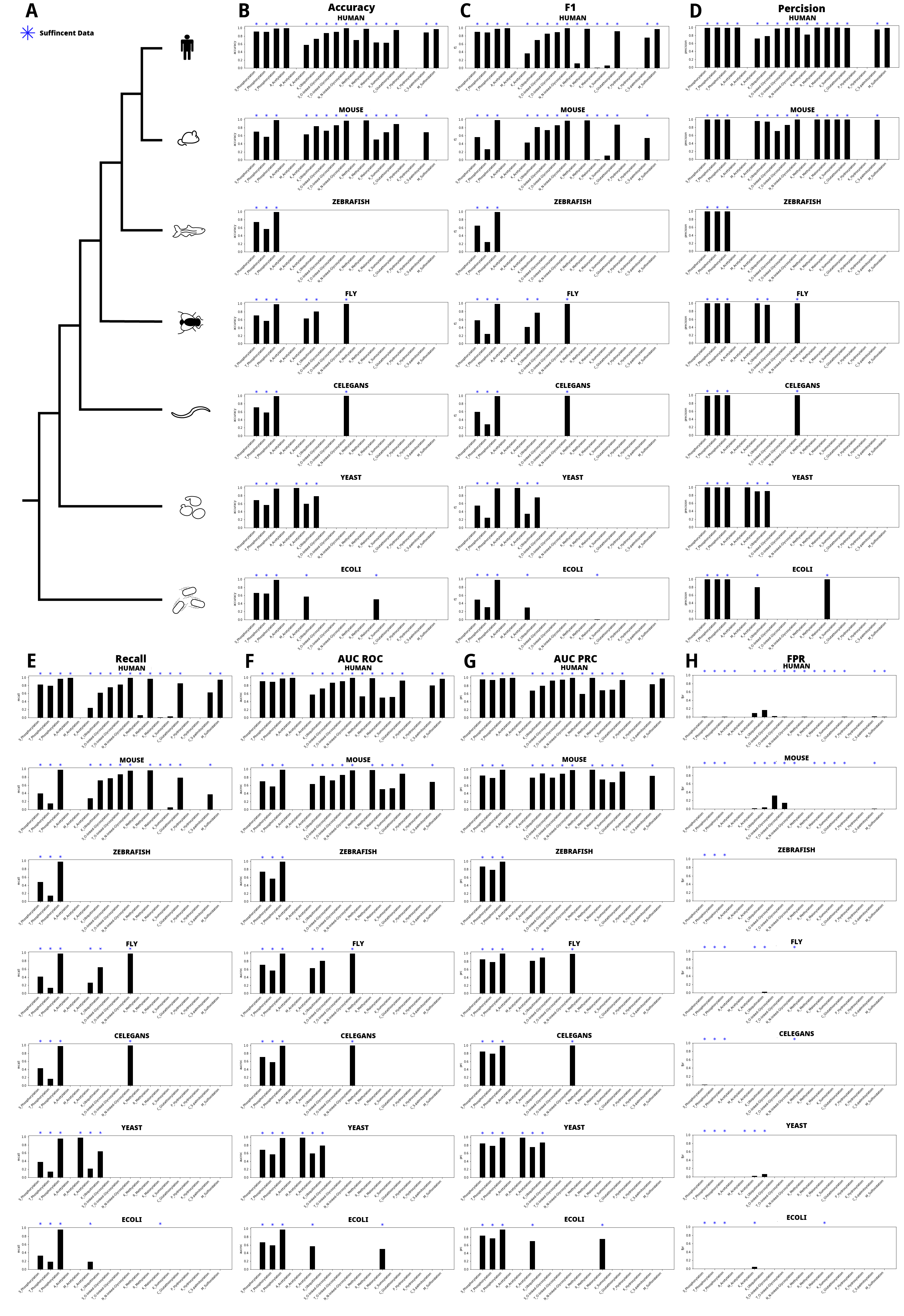
